# Morphological and Phylogenetic Resolution of *Diplodia corticola* and *D. quercivora*, Emerging Canker Pathogens of Oak (*Quercus* spp.), in the United States

**DOI:** 10.1101/2020.05.08.084111

**Authors:** Savannah L. Ferreira, Cameron M. Stauder, Danielle K.H. Martin, Matt T. Kasson

## Abstract

In Mediterranean Europe and the United States, oak species (*Quercus* spp.) have been in various states of decline for the past several decades. Several insect pests and pathogens contribute to this decline to varying degrees including *Phytophthora cinnamomi, Armillaria* spp., various insect defoliators, and additionally in the U.S., the oak wilt pathogen, *Bretziella fagacearum*. More recently, two emerging canker pathogens, *Diplodia corticola* (*Dc*) and *Diplodia quercivora* (*Dq*) have been implicated in causing dieback and mortality of oak species in Europe and in several regions in the United States. In 2019, a fungal survey was conducted in the Mid-Atlantic region of the Eastern U.S., including Maryland, Pennsylvania, Virginia, and West Virginia to determine the range and impact of *Dc* and *Dq* on forest health within the U.S. A total of 563 oak trees between red and white oak family members were evaluated across 33 forests spanning 18 counties. A total of 32 *Diplodia* isolates encompassing three *Diplodia* spp. were recovered from 5,335 total plugs collected from the 13 of 18 sampled counties. Recovered *Diplodia* species included *Dc, Dq*, and *D. sapinea* (*Ds*), as well as *Botryosphaeria dothidea* (*Bd*), a closely related canker pathogen in the *Botryosphaeriaceae*. Both *Dc* and *Ds* were recovered from red and white oak family members, whereas *Dq* was exclusive to white oak family members and *Bd* to red oak family members. Of these species, *Dc* was most frequently isolated followed by *Dq, Ds*, and *Bd*. Overall, mortality was relatively low across all sampled counties, indicating that these fungi, at the levels that were detected, are not widely inciting oak decline across the region, but more likely are acting opportunistically when the environment is conducive for disease. In an attempt to better understand the relationships among *Dc* and potentially their geographic origin(s), a multi-gene phylogenetic study and corresponding morphological study were conducted. A total of 49 *Diplodia* isolates from Spain, France, Italy, and the U.S. were assessed. Across all isolates and geographic regions, *Dc* formed a strongly supported monophyletic clade sister to *Dq* and included two strongly supported subclades, one that included isolates from Spain and California and a second that included isolates from Italy, Maryland, and West Virginia. Both subclades also exhibited overlapping spore measurements. These results support *Dc* as a cosmopolitan pathogen, native to both Europe and the U.S. with the possibility of secondary introductions.

## Introduction

Oak species (*Quercus* spp.), common to mixed hardwood forests of the Mid-Atlantic region of the eastern U.S., have been in various states of decline over the past several decades (Oak et al. 2016). Several pests and pathogens have been found to contribute to varying degrees, including *Bretziella fagacearum* (Bretz) Z.W. de Beer, Marinc., T.A. Duong & M.J. Wingf. (formerly *Ceratocystis fagacearum*), the causal agent of oak wilt (MacDonald et al. 2001, Jacobi and MacDonald 1980, Amos and True 1967, Shigo, A.L., 1958), *Phytophthora cinnamomi* Rands, the causal agent of *Phytophthora* root rot (McConnell and Balci 2014), and various insect defoliators, including gypsy moth (*Lymantria dispar* L.) (Tiberi *et al*. 2016, Asaro and Chamberlin 2015).

The symptomatology of these declines includes combinations of acute wilt, blighted leaves, external cankers, necrotic lesions, severe dieback, and in advanced stages, mortality (Reed et al. 2018, Smith and Santosz 2018, Martin et al. 2017, Munck et al. 2017, Martin and Munck 2017, Aćimović et al. 2016, Dreaden et al. 2011, Lynch et al. 2010). Abiotic stresses, including drought, flooding, competition, and mechanical damage, have also contributed to reduced vigor in native oak forests, which may predispose trees to infection by opportunistic pathogens (Bostock et al. 2014, Schomaker et al. 2007).

More recently, *Diplodia* spp., a genus associated with “bot” canker diseases of various angiosperm hosts, have been implicated in causing disease in red and white oak family members across the U.S. This follows closely behind detections of *Diplodia corticola* A.J.L. Phillips, A. Alves & J. Luque across Europe in association with regional oak decline in Spain, Portugal, France and Italy (Linaldeddu et al. 2017, Lazzizera et al. 2008, Alves et al. 2004).

In fall 2018, sampling for *Diplodia* spp. was conducted in areas with reported red oak (*Q. rubra* L.) dieback and blight in WV and MD. Fungal isolation from symptomatic tissues confirmed *Diplodia corticola* (*Dc*) in several sites in WV. Additionally, *Diplodia quercivora* (*Dq*) was identified from *Q. montana* Willd. at one site in MD, providing the first confirmation of *Dq* causing disease on a white oak family member in the Mid-Atlantic region (Haines et al. 2019). Prior to this discovery, *Diplodia quercivora* had been reported on southern live oak (*Q. virginiana* Mill.), another white oak family member, in FL (Dreaden et al. 2014). The apparent lack of host specificity and broad geographic distribution of *Dc* on both red and white oak family members informed our initial hypothesis that *Dc* was a generalist cosmopolitan pathogen while *Dq* has a more limited geographic range and is restricted to white oak family members.

These findings also raised important questions regarding the possible contribution by *Diplodia* spp. to the larger decline patterns across the landscape that had been casually observed but inadequately assessed, prompting a closer look at the USDA Forest Service’s Forest Inventory Analysis (FIA) data. To assess oak decline in the Mid-Atlantic region, we used the EVALIDator Web application to delimit annual mortality rates of red oaks by county (annual mortality volume/live volume) (data not shown). This analysis provided county-level resolution of mortality and identified 16 counties across DE, NJ, PA, VA, WV with between 5-17.4% annual mortality with the remaining counties reporting < 5% mortality.

To better understand the potential contribution by *Diplodia* spp. to oak decline occurring in the Mid-Atlantic region, we investigated the geographic distribution and impact of *Dc* and *Dq* on native oak species. The objectives of this study included: 1) Determine the incidence of *Diplodia* spp. in counties with varying levels of mortality in order to examine the correlation between disease severity and overall decline; and 2) Determine phylogenetic relationships among collected *Diplodia* isolates collected in the Mid-Atlantic region and isolates from allopatric populations in Europe and elsewhere in the U.S. Describing these relationships using phylogenetic inference as well as assessing diagnostically informative patterns among morphological measurements was important to investigate the nativity of these fungi as evidence supporting a cosmopolitan distribution is limited. Together, the results of this study offer insight into the ecology of *Diplodia* spp. and their possible contribution to the wide-spread oak decline in the Mid-Atlantic region.

## Materials and Methods

### Site selection

Counties were randomly selected for sampling based on red oak mortality percentages generated using USDA Forest Service FIA data within the EVALIDator Web application. The available FIA data separated county-level oak mortality into four distinct levels: < 1 % mortality (blue), 1.1─2.5% mortality (green), 2.6─5% mortality (yellow) and 5.1─17.4% mortality (red). Using a random number generator, one county from each mortality level was selected in MD, PA, VA, and WV (Figure 1).

**Figure 1.**
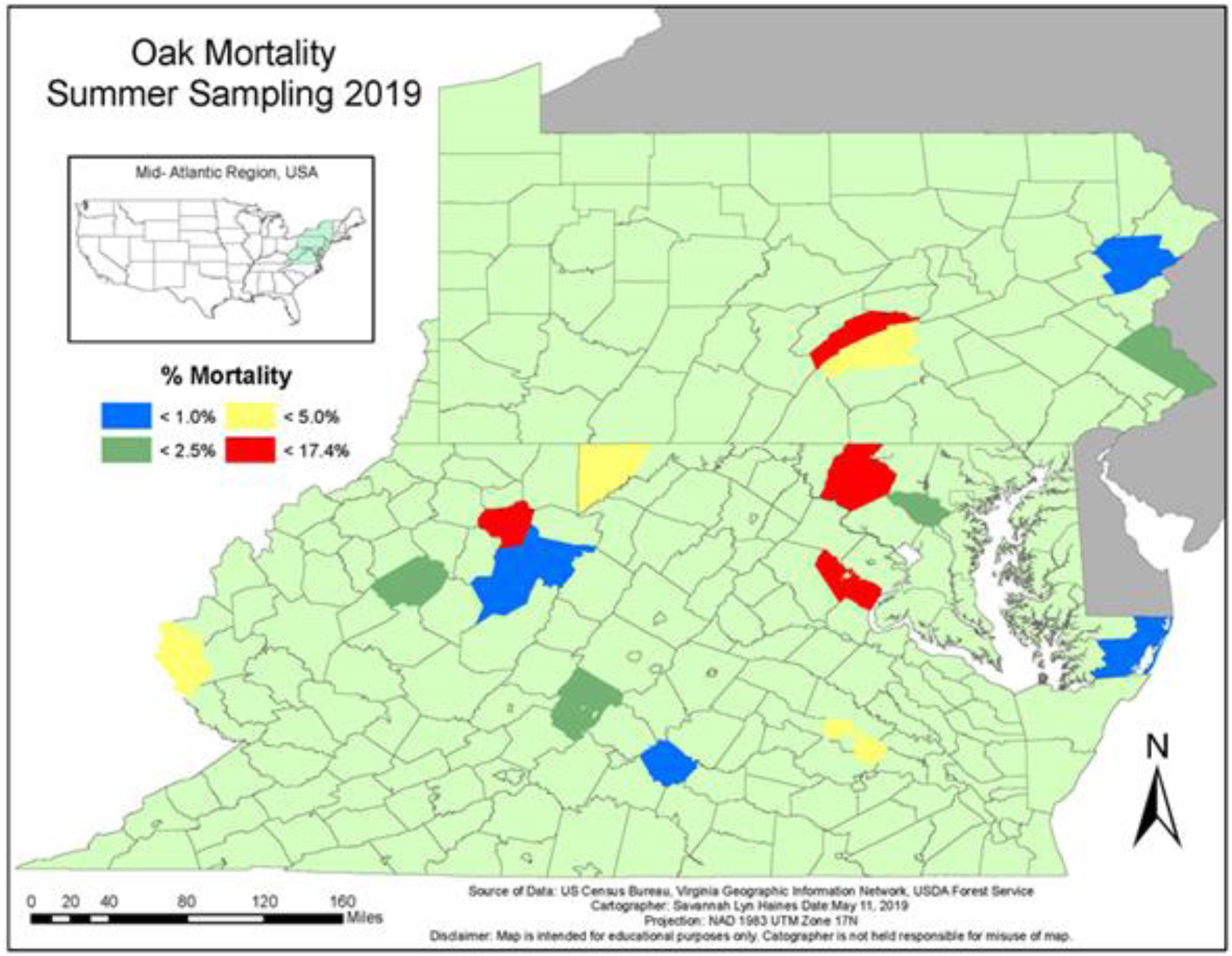
County-level sampling locations. Colored counties represent each % Mortality level of interest (blue: < 1 % mortality, green: 1.1─2.5% mortality, yellow: 2.6─5% mortality and red: 5─17.4% mortality in the four selected states (MD, PA, WV, VA)) in the lower Mid-Atlantic Region.

Two geographically separated forests in each of the selected counties were chosen based on oak presence, permissibility, accessibility, and ownership. Counties that did not have two forests that could meet that criteria were still included, but 30 trees were sampled in one forest, instead of the target of 15 trees per forest. These efforts resulted in the sampling of 31 forests in 16 counties (Supplemental Table 1). In addition to the counties stratified by mortality class, Green Ridge State Forest in Allegany County, MD and Seneca State Forest in Pocahontas County, WV were included to follow-up on the previous first detections of *Dq* in MD (Haines et al. 2019) and *Dc* in WV (Martin et al. 2017). Using publicly available forest boundary shapefiles, ArcGIS random plot generator created 30 labeled geographical coordinates (plots) per forest within each of the 33 forests. Plots in an individual forest were selected by their assigned number and their accessibility. Lower numbered plots were prioritized however, due to the randomized nature of this design, some lower number plots were not accessible and had to be skipped.

### Field sampling

At each forest, the randomly generated plot coordinates were used to establish the center of each variable radius plot. From the plot center, a 10 basal area factor prism was used to identify which trees were categorized as “in trees”, then tree species identity, diameter at breast height (DBH), and crown class were recorded. For all recorded oak trees, dieback rating (Figure 2) and the number of outwardly symptomatic tissues (cankers and necrotic “sooty” lesions) (Figure 3) were also recorded. Crown dieback ratings consisted of 5 categories: 1: 0-24% dieback, 2: 25-49% dieback, 3: 50-74% dieback, 4: 75-99% dieback and 5: 100% dieback (dead). Symptomatic tissues were counted and categorized into 6 classes: 0= no outward symptoms, 1= 1-3 visible cankers, 2= 4-6 visible cankers, 3= 7-9 visible cankers, 4= more than 9 visible cankers, NA= not applicable because it is dead. All observed dieback ratings were compared between oak family and outward symptoms classes using chi-square’s likelihood-ratio statistic.

**Figure 2.**
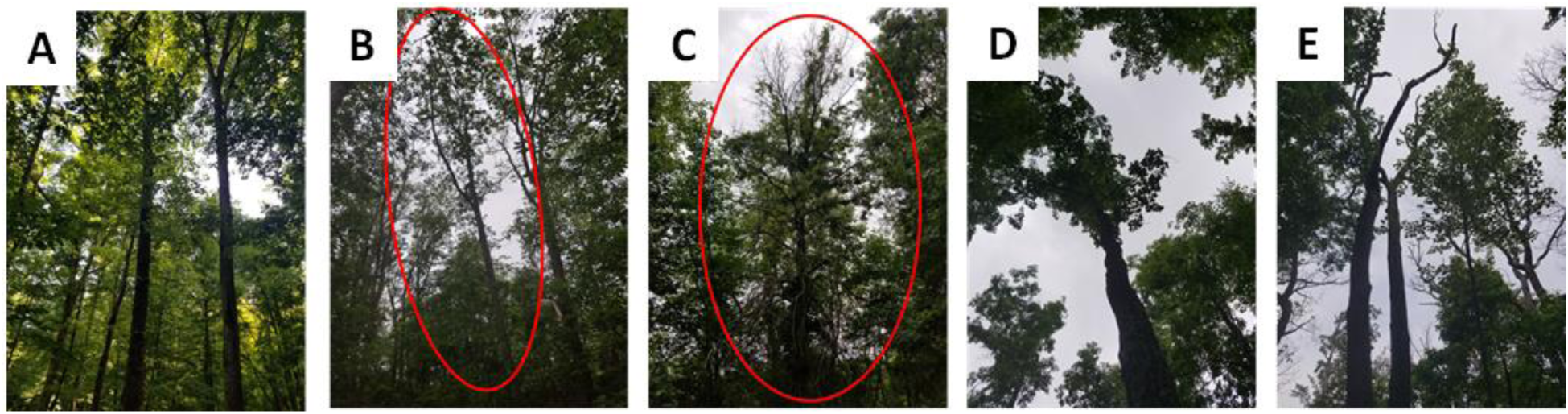
Progression of dieback in sampled oaks: A) 0-24%, B) 25-49%, C) 50-74%, D) 75-99%, E) 100%.

**Figure 3.**
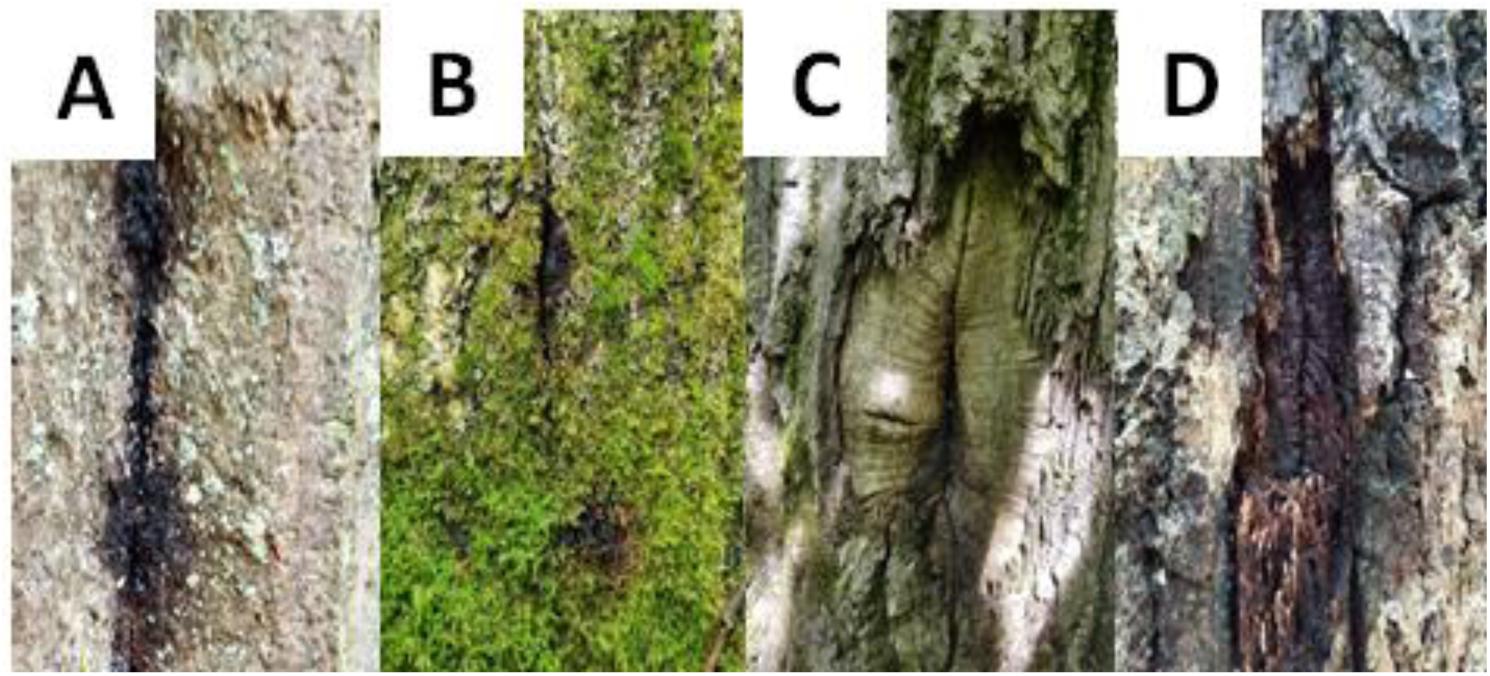
Examples of symptomatic tissue sampled for *Diplodia* spp. infection. A-B) Sooty lesions C-D) External cankers.

Observed oak trees were micro-sampled on the main bole with a sterile bone marrow biopsy tool; a small 0.5 −1 mm diameter tool that penetrates the tree (approx. 1-2 mm) to excise outer bark tissues to the vascular cambium (Stauder et al. 2019). A total of eight plugs were sampled per asymptomatic tree; four at DBH and four at approximately soil level in each of the cardinal directions. If the tree is outwardly symptomatic with sooty lesions and/or cankers (Figure 3), an additional eight bark sample plugs were taken in and around the symptomatic tissue. The extractor was flame-sterilized between symptomatic tissue and asymptomatic tissue on a singular tree as well as between trees. This resulted in a maximum of 16 bark sample plugs being collected per tree, with 15 trees being sampled per forest. *Diplodia* presence and *Diplodia* species obtained were compared to oak families using chi-square’s likelihood-ratio statistic.

If the prism plot did not yield 15 oak trees, the next plot that met accessibility and numeric criteria would be sampled, until 15 oak trees in the forest were micro-sampled. In total, 30 oak trees were sampled evenly across two forests per county with two exceptions. Due to access limitations in Beech Fork Wildlife Management Area in Wayne County, WV and the Monongahela National Forest in Randolph County, WV, 13 trees and seven trees were micro-sampled, respectively, resulting in Wayne County having 30 total sampled trees and Randolph county having only 27 sampled trees. All observed outward symptoms were compared between oak family and dieback ratings using chi-square’s likelihood-ratio statistic.

### Sample processing and fungal isolation

Field-collected bark sample plugs were surface disinfested in a 1:9 commercial bleach: water solution for 14 minutes and plated on glucose yeast extract agar with antibiotics (GYEA) for one week to observe fungal growth. Fungal isolates growing from the plugs were identified based on morphology and included *Penicillium, Trichoderma*, and *Pestalotiopsis* species. All *Diplodia* growing from the plugs were subcultured and retained as pure cultures. After seven days, any plugs that did not have fungal colonization were counted and classified as “uncolonized”. If the percentage of these uncolonized plugs exceeded 40% of the total plugs, plates were maintained for an additional week and previously uncolonized plugs were reassessed as described above.

### Isolate identification and phylogenetic analysis

In addition to *Diplodia* isolates retained from the field, 23 additional *Diplodia* spp. isolates were acquired from various university and government collections in Europe and the U.S. (Table 1). These additional isolates were isolated from *Q. alba, Q. agrifolia, Q. cerris, Q ilex, Q. montana Q. rubra, Q. suber, Q. velutina*, and *Q. virginiana* from France, Italy, Spain, and the U.S.

**Table 1:** Strain table for previously collected *Diplodia* spp. from various university and government collections used for phylogenetic and morphological analysis.

| Isolate ID | Other ID | Host | Location | *Diplodia* sp.<sup>z</sup> | GenBank Accession Numbers |  |  | Collector |
| --- | --- | --- | --- | --- | --- | --- | --- | --- |
|  |  |  |  |  | ITS | EF-1α | LSU |  |
| CJL155 | CBS 112071 | *Q. ilex* | Spain | *Dc* | --- | --- | --- | J. Luque |
| CJL245 | CBS 112072 | *Q. ilex* | Spain | *Dc* | --- | --- | --- | J. Luque |
| CJL248 | CBS 112073 | *Q. suber* | Spain | *Dc* | AY268420 | TBD | TBD | J. Luque |
| CJL255 | - | *Q. ilex* | Spain | *Dc* | TBD | TBD | TBD | J. Luque |
| CJL338 | CBS 112547 | *Q. ilex* | Spain | *Dc* | --- | --- | --- | J. Luque |
| CJL364 | - | *Q. cerris* | Italy | *Dc* | TBD | TBD | TBD | J. Luque |
| CJL47 | CBS 678.88 | *Q. suber* | Spain | *Dc* | --- | --- | --- | J. Luque |
| DC103 | QURU2 O 10 | *Q. rubra* | WV, USA | *Dc* | TBD | TBD | TBD | D. Martin |
| DC105 | QURU2 R | *Q. rubra* | WV, USA | *Dc* | TBD | TBD | TBD | D. Martin |
| DC110 | B012 | *Q. rubra* | MA, USA | *Dc* | --- | --- | --- | I. Munk |
| DC113 | HR 25-1-1 | *Q. rubra* | MA, USA | *Dc* | TBD | TBD | TBD | S. Aćimović |
| DC117 | OTL18-1 | *Q. rubra* | ME, USA | *Dc* | TBD | TBD | TBD | S. Aćimović |
| DC118 | OTCT3-1 | *Q. rubra* | ME, USA | *Dc* | TBD | TBD | TBD | S. Aćimović |
| PL1477 | MTF11 AB | *Q. virginiana* | FL, USA | *Dc* | --- | --- | --- | J. Smith |
| PL1480 | SVH1 AB | *Q. virginiana* | FL, USA | *Dc* | --- | --- | --- | J. Smith |
| PL1518 | OSC3 B1 AB | *Q. virginiana* | FL, USA | *Dc* | TBD | TBD | TBD | J. Smith |
| UCR1246 | JN693502 | *Q. agrifolia* | CA, USA | *Dc* | JN693502 | JQ512108 | TBD | S. Lynch |
| UCR1339 | JQ411401 | *Q. agrifolia* | CA, USA | *Dc* | TBD | TBD | TBD | S. Lynch |
| UCR488 | JN693501 | *Q. agrifolia* | CA, USA | *Dc* | JN693501 | JQ512132 | TBD | S. Lynch |
| WI 03-35 | - | *Q. alba* | WI, USA | *Dc* | TBD | TBD | TBD | G. stanosz |
| CJL374 | - | *Q. suber* | France | *Dm* | TBD | TBD | TBD | ? |
| DC114 | WHG PRKL-1-5 | *Q. rubra* | MA, USA | *Dq* | TBD | TBD | TBD | S. Aćimović |
| DQ703 | MK160499 | *Q. montana* | MD, USA | *Dq* | TBD | TBD | TBD | S. Ferreira |
| MFLUCC 100098 | - | - | - | *Bf* | JX646789 | JX646854 | JX646806 | B. Slippers |

<sup>z</sup> *Dc*, *D. corticola*; *Dq*, *D. quercivora*; *Dm*, *D. mutila*, *Bf*, *Botryosphaeria fusispora* (outgroup)

All *Diplodia* isolates were transferred from stock culture to a GYE agar plate that had a surface layer of sterile cellophane and grew for approximately one week before being harvested for DNA extraction using a Wizard® kit (Promega, Madison, WI, USA) and resuspended in 75 µl Elution Buffer (Alfa Aesar, Ward Hill, MA, USA). For each *Diplodia* isolate, PCR amplification and sequencing of the internal transcribed spacer region (ITS) using primers ITS4 and ITS5 (White et al., 1990), elongation factor gene (EF-1α) using primers EF1728F (Carbone and Kohn, 1999) and EF1-1567R (Rehner 2001) and the large subunit (LSU) using primers LR5 (Vilgalys and Hester, 1990) and LROR (Rehner and Samuels, 1994) were conducted.

To amplify these products, 10 µl of molecular-grade water (G-Biosciences, St. Louis, MO, USA), 12.5 µl of Master Mix (Bioline, London, UK), 1 µl of each respective forward and reverse primer (IDT, Coralville, IA, USA), and 1.5 µl of template DNA were combined. ITS and EF-1α gene regions were amplified using a three-stage PCR reaction: 95 °C for two minutes, followed by 35 cycles of 95 °C for 30 seconds, 56 °C for 30 seconds, 72 °C for 60 seconds with a final extension at 72 °C for seven minutes. LSU gene region amplification was as follows: 95 °C for two minutes, followed by 35 cycles of 95 °C for 30 seconds, 51.1 °C for 45 seconds, and 72 °C for 90 seconds with a final extension at 72 °C for five minutes. PCR products were prepared for sequencing using ExoSAP-IT (Affymetrix, Santa Clara, CA, USA) according to the manufacturer’s recommendations and then Sanger-sequenced (Eurofins, Louisville, KY, USA) using their respective PCR primers.

All sequences for the following phylogenetic analyses were clipped in CodonCode Aligner v. 5.1.5 and corrected for missing or miscalled nucleotides by referencing chromatograms. Maximum-likelihood and Bayesian inference analyses were conducted for single gene and concatenated three-gene (ITS, EF-1α, LSU) sequences. Additional reference sequences were selected from those available NCBI Genbank (Table 1). Each of the three analyzed gene regions were separately aligned using MAFFT (Katoh and Stanley, 2013) on the Guidance 2.0 server (Landan and Graur, 2008; Sela et al., 2015), and all residues having Guidance scores < 0.5 were masked (Macias et al., 2020). Aligned sequences were then concatenated into a single three-gene sequence using FaBox (Villesen 2007). For maximum-likelihood analyses, the substitution model was selected from Model Test AICc scores in MEGA v10.1.7 (Stecher et al., 2020), and 1,000 bootstrap replicates were used for the analysis. Bayesian inference (BI) analyses were completed using MrBayes v. 3.2.7 (Ronquist et al., 2012). The best fit nucleotide selection model and the rate of substitution were selected by MrBayes. BI runs were stopped once the standard deviation of split frequencies fell below 0.01 and then checked for convergence in Tracer v. 1.7.1 (Rambaut et al., 2018). Trees were formatted for publication using FigTree v. 1.4.4 (Rambaut, 2017), Adobe Illustrator v. 24.1, and Inkscape v. 0.92.5. All trees represent the ML topology with both ML and BI values at branches (ML/BI). Strong boot-strap support is represented by ML values ≥ 70% and moderate support if boot-strap values were between 50% and 69%.

### Spore production and associated measurements

*Diplodia spp*. cultures were grown on 1/10th strength GYE to induce pycnidia production. Selected isolates were transferred as 0.5 cm fully colonized plugs to 1/10th GYE and left under continuous light for four weeks (Figure 4). After one-month, morphological observations including the presence of pycnidia and spore measurements were made. Recovered spore masses from pycnidia were mounted on slides containing lactophenol to observe spore size and morphology. The length and width of twenty-five randomly selected spores were measured at the longest point using a Nikon Eclipse E600 microscope and NIS-Elements BR 3.2 under 40X magnification. Spore sizes and associated features (i.e. number of septations and spore pigmentation) were compared between species and among geographic locations using analysis of variance (ANOVA) and then between locations using Tukey Kramer’s post hoc test.

**Figure 4.**
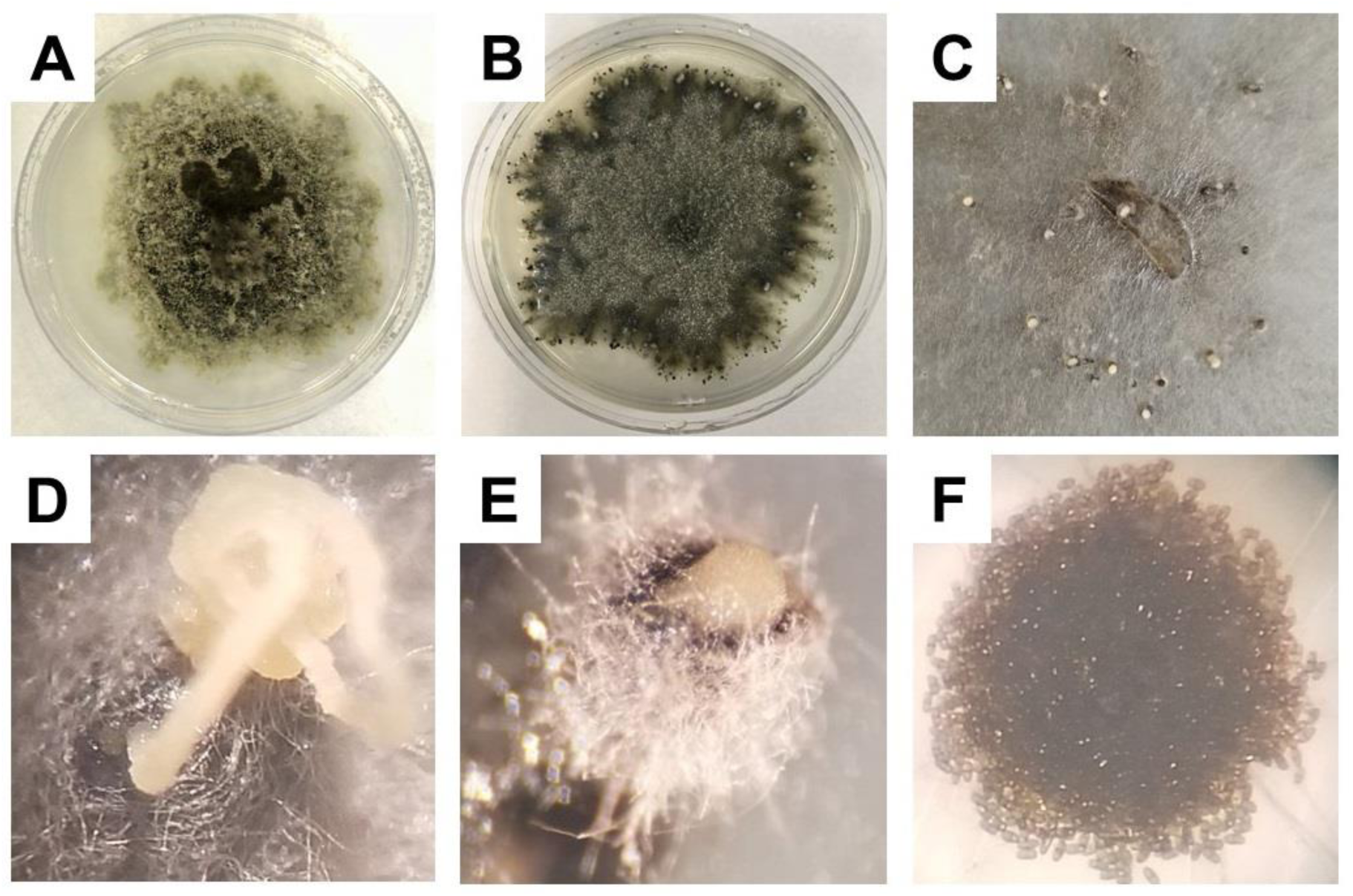
Representative *Diplodia* cultures exhibiting pycnidia production: A) *Dc* isolate growing on GYEA, B) *Dc* isolate developing pycnidia precursors on 1/10^th^ GYE, C) *Dc* isolate exuding conidia from numerous pycnidia on 1/10^th^ GYE, D) Conidial tendril being exuded from pycnidia, E) Mass of conidia being exuded from pycnidia, F) Mass of brown septate conidia.

## Results

A total of 5,335 bark sample plugs were processed from 563 oak trees spanning 33 individual forests across the four sampled states. Approximately 13% (688 samples) of the collected bark samples failed to yield any fungal isolates. Of the remaining 4,647 samples that yielded fungal growth, approximately 1% (32 samples) yielded isolates of *Diplodia* spp. representing 13 of the 18 sampled counties (Figure 5). The remaining (4,615) bark samples yielded various environmental fungi, including *Trichoderma* spp., *Penicillium* spp. and *Pestalotiopsis* spp., although these fungi were not further characterized since the primary focus was delimitation of *Diplodia* spp.

**Figure 5.**
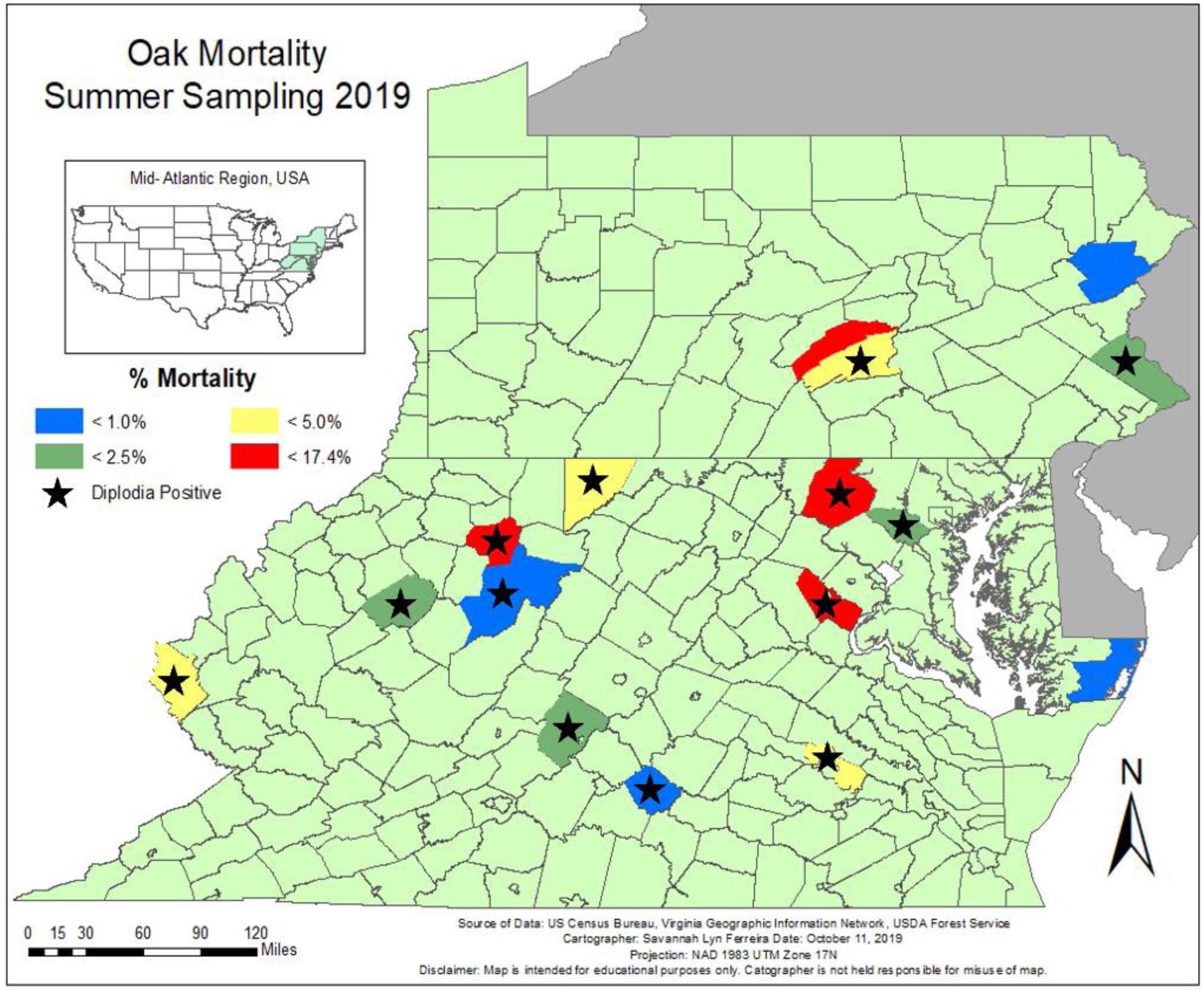
Oak mortality sampling map showing *Diplodia* positive counties (n = 13). Sampling locations included the following counties: MD: Worcester (blue), Howard (green), Garrett (yellow) and Fredrick (red); PA: Monroe (blue), Bucks (green), Perry (yellow) and Juniata (red); VA: Appomattox (blue), Rockbridge (green), Henrico (yellow) and Prince William (red); WV: Randolph (blue), Braxton (green), Wayne (yellow) and Barbour (red).

ITS barcoding confirmed the identities of the 32 isolates of *Diplodia* spp. retained from the field survey samples with 21 *D. corticola* (*Dc*) isolates (96-100% ITS sequence similarity to *Dc* accessions available on NCBI Genbank), six *D. quercivora* (*Dq*) isolates (94-100% ITS sequence similarity to *Dq* accessions available on NCBI Genbank), three *D. sapinea* (*Ds*) isolates (97-100% ITS sequence similarity to *Ds* accessions available on NCBI Genbank), and two *Botryosphaeria dothidea* (*Bd*) isolates 94-99% ITS sequence similarity to *Bd* accessions available on NCBI Genbank). For details of specific locations and isolates recovered see Supplemental Table 2.

### Field sampling

Five hundred sixty-three oak trees were sampled between red (n = 191) and white oak (n = 372) family members in 33 forests in the Mid-Atlantic region, 507 of which were sampled for fungal colonization. The remaining 56 trees were excluded from fungal sampling due to sample size restrictions as written in landowner sampling permits. The red oak family members observed included red oak (*Q. rubra*) (n = 146), pin oak (*Q. palustris* Münchh.) (n = 16) and turkey oak (*Q. laevis* Walter) (n = 29), while white oak family members included white oak (*Q. alba* L.) (n = 194), chestnut oak (*Q. montana*) (n = 167), southern white oak (*Q. bicolor* Willd.) (n = 10) and willow oak (*Q. phellos* L.) (n = 1).

### Dieback

To assess plot-level oak decline, dieback for all oak trees that fell within the 10 basal area factor variable radius plot were evaluated and recorded. These included 191 ratings for red oak family members and 372 ratings for white oak family members. Observations for individual trees were grouped into 5 discrete categories, 1: 0-24% dieback, 2: 25-49% dieback, 3: 50-74% dieback, 4: 75-99% dieback and 5: 100% dieback (dead).

Oak family was a significant predictor of percent dieback (χ^2^= 16.10, df = 559; P= 0.0029), with white oak family members having higher counts for more severe dieback ratings than red oak family members (Table 2). Four hundred and three of the 563 observed trees had minimal dieback at a rating of 1, with 282 being white oak family members and 121 were red oak family members (Table 2). The remaining 160 total red and white oak trees measured had more severe dieback ratings of 2-5, with the majority (17.58%) with a rating of 2.

**Table 2:** Summary statistics for dieback rating by family for all observed oak trees. Data are partitioned by dieback categories: 1, 0-24% dieback; 2, 25-49% dieback; 3, 50-74% dieback; 4, 75-99% dieback; and 5, 100% dieback (dead) and oak family (RO = red oak and WO= white oak family members.

| **Oak Family** | **Dieback Category** |  |  |  |  | **Total** |
| --- | --- | --- | --- | --- | --- | --- |
|  | **1** | **2** | **3** | **4** | **5** |  |
| RO | 121<br>(21.5%) | 51<br>(9.1%) | 7<br>(1.2%) | 3<br>(0.5%) | 9<br>(1.6%) | 191<br>(33.9%) |
| WO | 282<br>(50.1%) | 52<br>(8.5%) | 13<br>(2.3%) | 7<br>(1.2%) | 22<br>(3.9%) | 372<br>(66.1%) |
| Combined | 403<br>(71.6%) | 53<br>(17.6%) | 20<br>(3.6%) | 10<br>(1.8%) | 31<br>(5.5%) | 563<br>(100%) |

<sup>z</sup> Counts (% incidence)

### Outward symptoms

As trees with severe dieback may express additional outward symptoms (e.g. sooty lesions, cankers), we compared dieback ratings to symptom classes. When excluding dieback as an outward symptom, 160 observed trees had at least one discrete symptom while 403 were outwardly asymptomatic or too dead to determine/assess symptoms. For symptomatic trees, the majority of observations across all symptom classes had a dieback rating of 1, followed by a dieback rating of 2 for symptom classes 1-3. A majority of trees in symptom class 4, excluding dieback category 1, had a dieback rating of 5 (Table 3).

**Table 3:** Summary statistics for dieback rating by symptom class for all observed oak trees. Data are separated by dieback category: 1, 0-24% dieback; 2, 25-49% dieback; 3, 50-74% dieback; 4, 75-99% dieback; and 5, 100% dieback (dead) and symptom class: 0, no outward symptoms; 1, 1-3 visible cankers; 2, 4-6 visible cankers; 3, 7-9 visible cankers; 4, > 9 visible cankers; and NA=, not applicable tree dead.

| Symptom<br>Class | Dieback Category |  |  |  |  | Total |
| --- | --- | --- | --- | --- | --- | --- |
|  | 1 | 2 | 3 | 4 | 5 |  |
| 0 | 294<br>(52.2%)<sup>z</sup> | 62<br>(11%) | 15<br>(2.7%) | 7<br>(1.2%) | 13<br>(2.3%) | 391<br>(69.4%) |
| 1 | 86<br>(15.3%) | 29<br>(5.2%) | 3<br>(0.5%) | 3<br>(0.5%) | 3<br>(0.5%) | 124<br>(22.1%) |
| 2 | 16 (2.8%) | 7<br>(1.2%) | 2<br>(0.4%) | 0 | 1<br>(0.2%) | 26<br>(4.6%) |
| 3 | 2<br>(0.4%) | 1<br>(0.2%) | 0 | 0 | 0 | 3 (0.5%) |
| 4 | 5<br>(0.9%) | 0 | 0 | 0 | 2<br>(0.4%) | 7 (1.3%) |
| NA | 0 | 0 | 0 | 0 | 12<br>(2.1%) | 12<br>(2.1%) |
| Total | 403<br>(71.6%) | 99<br>(17.6%) | 20<br>(3.6%) | 10<br>(1.8%) | 31<br>(5.5%) | 563<br>(100%) |

<sup>z</sup> Counts (% incidence)

Oak family was also a significant predictor of symptom class (χ^2^= 31.72, df = 559; P= 0.0001) with white oak family members being less likely to show conspicuous and discrete symptoms (e.g. cankers and sooty lesions) than red oak family members. Between both families, the majority of symptomatic trees fell into symptom class 1 (n = 124) (Table 4).

**Table 4:**
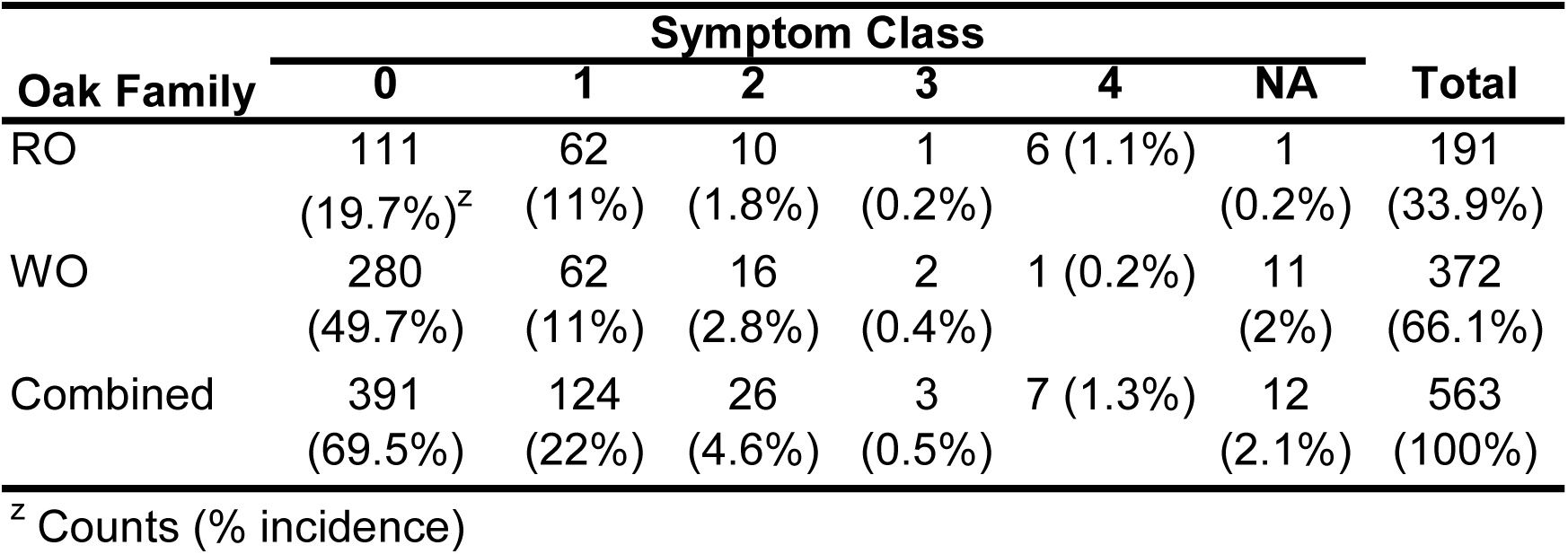
Summary statistics for symptom class by oak family. Data are separated by symptom class: 0, no outward symptoms; 1, 1-3 visible cankers; 2, 4-6 visible cankers; 3, 7-9 visible cankers; 4, > 9 visible cankers; and NA, not applicable dead tree and oak family: RO, red oak and WO, white oak family members.

### *Diplodia* presence

Five hundred and seven trees were sampled for *Diplodia* presence, which included 347 trees without any conspicuous and discrete symptoms and 160 with discrete symptoms (data not shown). Thirty-two isolates of *Diplodia* were isolated from 27 trees. From these trees, 13 *Diplodia* isolates were recovered from symptomatic bole tissues and 19 from outwardly asymptomatic bole tissues. Four trees had multiple isolates in either symptomatic or asymptomatic tissue, so only 28 isolates of *Diplodia* were used in the analysis.

Oak family was a significant predictor of *Diplodia* species (χ^2^= 17.48, df = 24; 0.0006). Sampled red oak family members yielded 60.71% (17 out of 32) of the recovered *Diplodia* isolates while white oak family members yielded 39.29% (11 out of 32) of the recovered isolates. The highest recovered *Diplodia* spp. was *Dc* (n = 18) with red oak family members having 15 isolates and white oak family members having three isolates. The second highest recovered species was *Dq* (n = 6) with all isolates recovered from white oak family members. The third most common isolate was *Ds* with three total isolates between both families, two from white oak and one from red oak family members. *Bd* was the least recovered species, with only one isolate from a red oak family member. Pathogenicity tests performed on 15 red oak seedlings failed to confirm *Bd* as a standalone pathogen of red oak, despite its recovery from symptomatic tissue (data not shown). Pathogenicity tests for *Ds* were not conducted due to time constraints.

### Phylogenetic studies

A total of 49 isolates were included in the single-gene and concatenated phylogenetic analyses. Three isolates served as either ingroup reference strains including *Dq* isolate DQ703 (Haines et al. 2019) and *Dc* isolate DC103 (Martin et al. 2017), or outgroup taxa, *Botryosphaeria fusispora* (Slippers et al. 2013) (Table 1). The three-gene concatenated phylogeny resolved all *Diplodia* species recovered into a monophyletic group with two major clades, one including *Dc* and *Dq* with a 95% bootstrap support and a second clade containing *Ds* with a 98% bootstrap support (Figure 6). The larger clade strongly supported *Dq* as a genealogically exclusive species with 81% bootstrap support. Among three single-gene trees, the four *Diplodia* species were genealogically exclusive in only the ITS tree.

**Figure 6.**
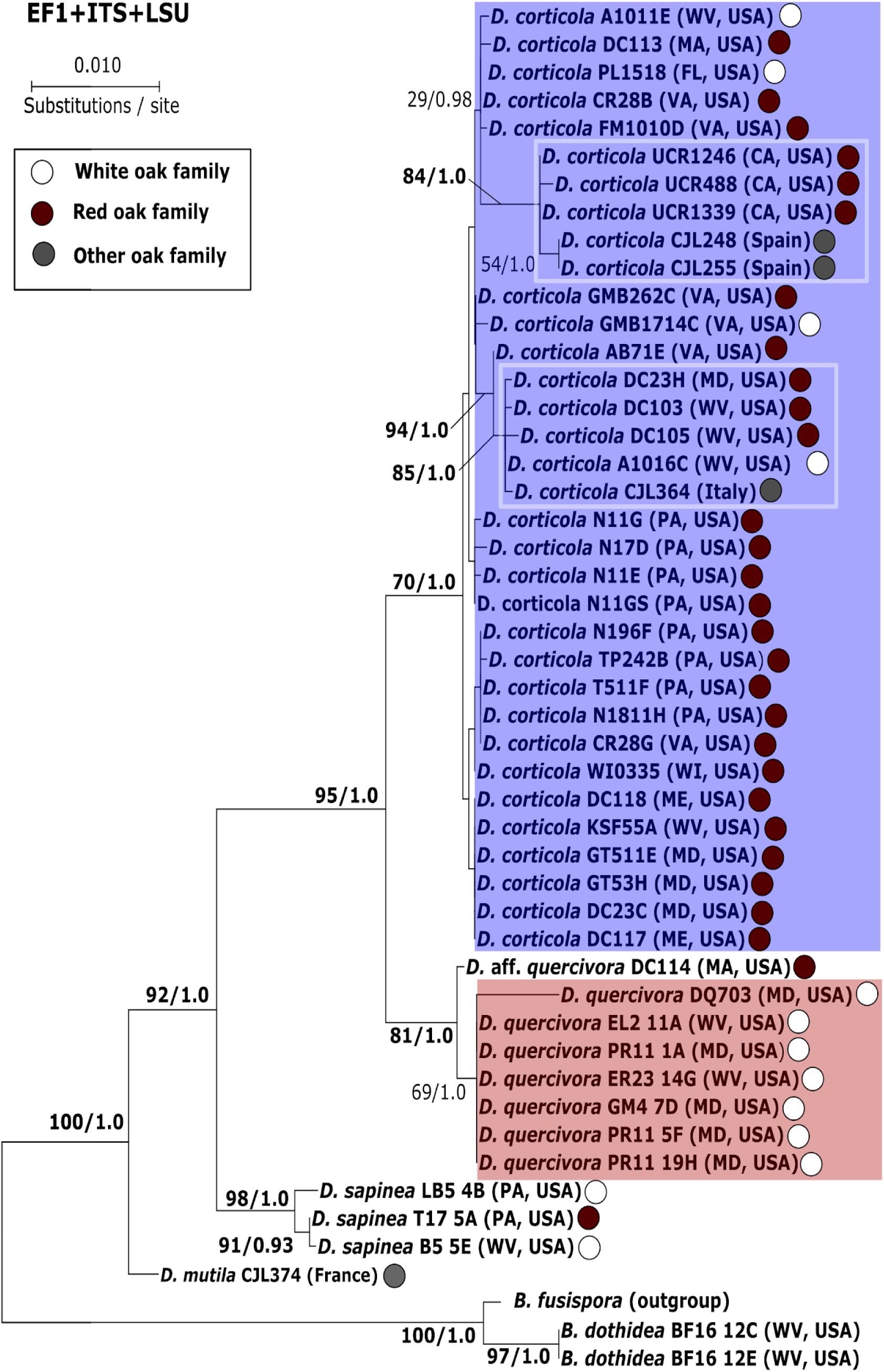
Concatenated phylogenetic tree for ITS, EF-1α and LSU gene regions showing relationships among *Diplodia* spp. and allied fungi. Topology and branch lengths shown are from the ML analysis. Bootstrap support and posterior probabilities are indicated near each node (ML/BI), and nodes with >50% support are labeled. White boxes indicate clades within *D. corticola* that were strongly supported.

### Spore production and associated measurements

Fifty-three isolates were plated on 1/10^th^ GYE under continuous light to induce sporulation; 30 from field survey and 23 from university and government collections. After four weeks, conidia were observed from 42 *Diplodia* spp. isolates. Pycnidia were typically visible within two weeks of plating, with spores becoming visible within the following week. Observed spore sizes were significantly different by species (*P* < 0.0001) and location (*P* < 0.0001).

*Dc* isolates had the second smallest spore size among *Diplodia* spp. observed, averaging 19.58 ± 1.71 µm (Figure 7). Among all isolates, *Dc* conidia were typically spherical to elliptical in shape and hyaline in color. Frequently, spores were observed as a light to medium brown with a single septation (Figure 8A). *Dc* analyzed spores from CA were the largest (21.02 ± 0.17 µm), followed by Spain (20.38 ± 0.14 µm), VA (20.18 ± 0.12 µm), MA (19.93 ± 0.21 µm), WV (19.68 ± 0.14 µm), MD (19.47 ± 0.15 µm), FL (19.42 ± 0.17 µm), ME (18.89 ± 0.21 µm), Italy (18.47 ± 0.30 µm), WI (18.37 ± 0.30 µm) and PA (18.34 ± 0.11 µm) (*P* < 0.0001).

**Figure 7.**
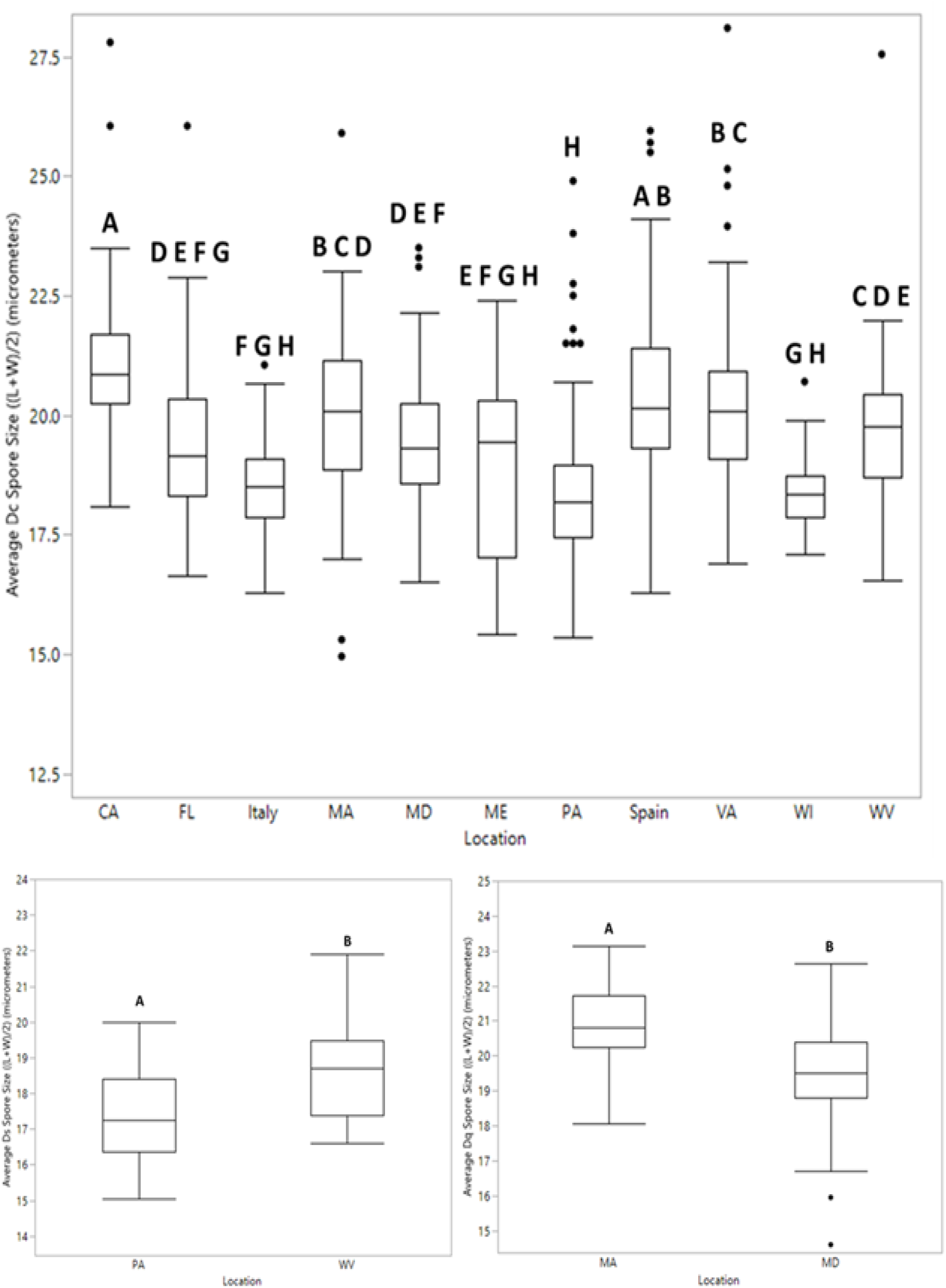
Box plots for average spore size ((Length + Width)/2) of *Diplodia* spp. Spore sizes compared between species and among geographic locations using ANOVA. Letter designation by Tukey Kramer’s ordered letter report shows significant differences by location. Single points outside the upper and lower extremes represent outliers and were included in analysis. A) *Dc* (N = 35) isolates by location (CA (n = 3), FL (n = 3), Italy (n = 1), MA (n = 2), MD (n = 4), PA (n = 7), Spain (n = 5), VA (n =6), WI (n = 1) and WV (n = 3). B) *Ds* isolates (N = 3) by location (PA (n = 2) and WV (n 1)). C) *Dq* (N = 4) isolates by location (MA (n = 1) and MD (n = 3)).

**Figure 8.**
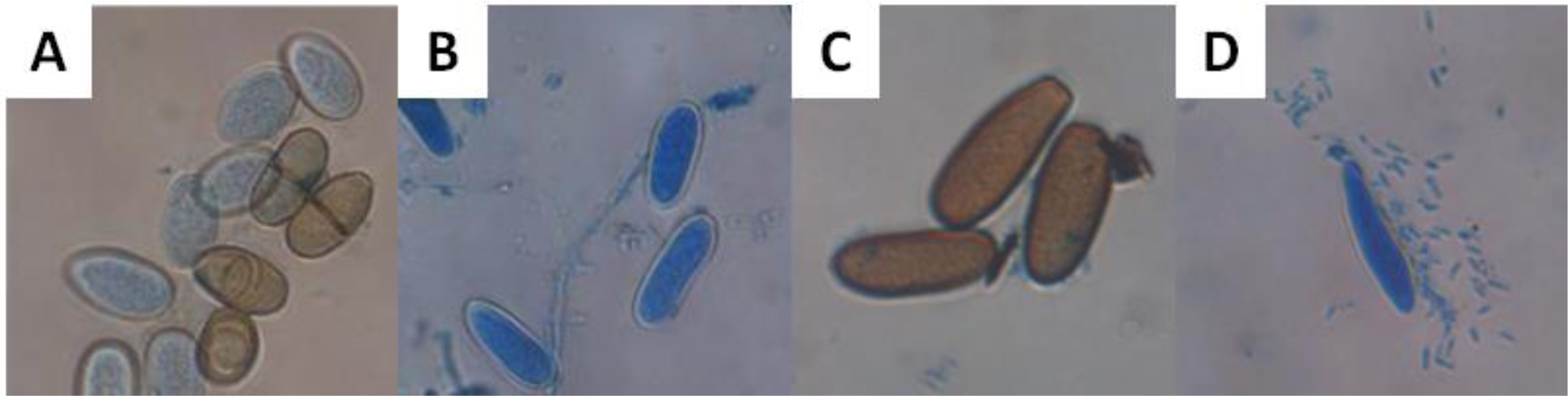
Representative conidia morphology of recoveredaps Diplodia spp. And allied fungi including A) *D. corticola*, B) *D. quercivora*, C) *D. sapinea*, and D) *Botryosphaeria dothidea*. Mountant is lactophenol + cotton blue. Spores are shown at 40X magnification, not to scale.

*Dq* had the second-largest observed spores, however, due to *Dq* rarely producing pycnidia on 1/10^th^ agar, only two out of six *Dq* isolates recovered during this study were analyzed. *Dq* spores averaged 19.79 ± 1.50 µm (Figure 7). These spores were hyaline in color and elliptical to oblong in shape (Figure 8B). These spores were not observed as being septate or pigmented. Spores from MA were significantly larger than spores from MD with measurements averaging 20.82 ± 0.27 µm and 19.45 ± 0.15 µm respectfully.

*Ds* isolates had the smallest spore size across all observed *Diplodia* species, averaging 17.81 ±1.38 µm. These spores were observed as elliptical to oblong in shape and brown in color with no observed septations (Figure 8C). For this species, spores from the WV were significantly larger (*P* < 0.0002) than the spores observed from PA which measured 18.61 ± 0.25 µm and 17.41 ± 0.18 µm, respectively (Figure 7).

The largest observed conidia were from the single recovered *Bd* isolate. This isolate also produced microconidia, but they were not measured in this study. The observed macroconidia averaged 25.69 ± 3.68 µm. The macroconidia were long elliptical in shape and hyaline to brown in color. Microconidia were also elliptical in shape and hyaline in color. No septations were observed in either micro or macroconidia (Figure 7D). Due to the observation of a single *Bd* isolate, spore sizes were not compared by location.

## Discussion

A regional survey was conducted in four states in the Mid-Atlantic region to observe the geographic distribution and level of damage caused by *Diplodia* spp. and explore their possible contributions to regional oak decline. The results of this study revealed that several species of *Diplodia* are present in the region, however, their individual presence and damage vary by location and hosts.

In MD and VA, *Dc* was recovered for the first time from red oak family members including both red oaks in MD and turkey oaks in VA. In WV and PA, although already present, *Dc* was recovered for the first time on white and chestnut oak in WV, and for the first time on pin oak in PA. Although typically a fungal pathogen of two- or three-needled pines (Brodde et al. 2018, Rajotte 2017), *Ds* was recovered in this study from both chestnut and northern red oaks in PA and represents the first occurrence in WV on white oak. Another fungal pathogen of numerous woody ornamental hosts (Turco et al. 2006, Sánchez et al. 2003), *Bd* was recovered for the first time from symptomatic red oak trees in WV. Isolates of *Dq*, which had been only recently confirmed from MD, were recovered for the first time on chestnut oaks in WV.

The low numbers of *Diplodia* obtained from this survey may not accurately portray the incidence or impact this fungal canker pathogen is having in the region. This low recovery could be the result of sub-optimal sampling weather, inefficient sampling methods, and/or due to the randomized nature of field locations. *Diplodia* may have also been underrepresented due to heterogeneous distribution of fungal mycelium within asymptomatic host tissues, and/or extracting plugs that were colonized by *Diplodia*-inhibiting fungi (Ferreira 2019). Using randomized sampling locations may have had the most drastic effect on recovered *Diplodia* spp. by potentially missing *Diplodia* positive trees. In both Seneca State Forest and Green Ridge State Forest, *Diplodia* was isolated and reported as being the causal agent of the observed decline (Haines et al. 2019, Martin et al. 2017). Upon resampling of those forests using the methods described in this paper, trees observed were in a separate geographic location in the forest and were *Diplodia* negative. Follow-up studies may provide insights into the size of outbreaks and abiotic and biotic factors that influence them.

*Diplodia*-positive trees were both outwardly asymptomatic and symptomatic, with symptomatic trees having dieback and/or bole symptoms. From the observations made in the field, symptomatic tissue was a significant predictor of dieback, however, they were not observed as large or frequent enough in this survey to cause large scale dieback or mortality. Based on our observations, white oak family members had fewer outward symptoms and had less isolated *Diplodia* spp. then the red oak family members. Due to oak decline’s history of other biotic complexes in addition to white oak family members’ symptom development, it is possible that *Diplodia* spp. have been overlooked as causal agents of other recent declines including Rapid White Oak Mortality (Reed et al. 2017). White oak family members may be responding faster to *Diplodia* infections due to the utilization of tyloses in compartmentalization, which acts as an additional response to wall off the vascular tissue (De Micco et al 2016, Meier 2008). This leads to the observation that when *Diplodia* is present, dieback is a better indicator of colonization in white oak family members, while outward symptom presence is a better indicator of colonization in red oak family members.

*Diplodia* spp. have been found in comparable numbers across 13 out of 18 counties and is present in all FIA mortality classes sampled. Although observed in a wide range of locations, this pathogen had low recovery across the region. When recovered isolates were observed with government collection, the results revealed that many of the *Dc* isolates examined were not phylogenetically distinct (with strong bootstrap support) based on location or the gene regions studied.

Previous studies indicate that *Dc* may have been inadvertently introduced into the U.S. from areas in Europe, where *Dc* has been associated with cork oak decline since the early 2000s (Lynch et al. 2013). The results of our phylogenetic analysis of the 3-gene combined dataset provide strong support for linkages between isolates from CA and Spain (84% bootstrap support) as well as between isolates from Italy, MD, and WV (85% bootstrap support). California and Spain are two regions that are among the world’s leading wine producers, both of which have a history of *Dc* causing disease on grapes (*Vitis* spp.) (Varela et al. 2011, Úrbez-Torres et al. 2010). This observation, along with similar individual spore sizes, provides evidence that some *Dc* populations causing disease in CA may have been introduced from Spain. Likewise, the strong support between Italy, MD, and WV may provide evidence of the potential for secondary introductions, although these possible linkages are less clear than CA-Spain.

Ultimately, cross-pathogenicity studies are needed to compare isolates from Europe and the U.S. to better understand the host specificity and virulence of strains. Similar studies of oak wilt in European versus American oaks in common orchards in WV revealed high susceptibility of European white oak family members compared to limited susceptibility of U.S. white oak family members (MacDonald et al. 2001).

Morphological studies have also proven useful in describing and comparing species. Although congruence between morphological trends and phylogenetic groupings have been shown for several fungi, the number of isolates used in this study were not sufficient nor consistent enough to make meaningful comparisons between sampled locations. Spain had the most observed isolates (n = 5), followed by CA (n = 3) and FL (n = 3), with fewer counts for the other observed isolates. Spain and CA isolates exhibited the largest spores across all locations examined, but larger trends based on these morphological features across *Dc* were not consistent or phylogenetically informative.

Although the specific strains of *Dc* recovered from CA align with isolates from Spain and may indicate an introduction from Europe, the other states within the U.S. affected by *Dc* are more geographically isolated from international ports. However, reported U.S. states are mostly coastal, which could mean that *Dc* is acting opportunistically in areas that are more sensitive to the changing climate, or that *Dc* is being introduced in numerous locations due to imports from Europe. States including FL, MD, VA had overlapping spore morphologies that were supported in the 3-gene concatenated tree, supporting the hypothesis that *Dc* is a native pathogen in the U.S. Similar European-U.S. groupings are apparent between Italy, MD and WV isolates in both the 3-gene tree (85% boot-strap) and spore morphologies.

The low incidence of isolation coupled with the geographic overlap of documented declines supports the observation that *Dc* is contributing to oak decline in the U.S., however, it is not likely the inciting agent of widespread mortality.

## Conclusion

*Diplodia corticola* has been implicated over the last couple decades in contributing to oak decline in Mediterranean Europe and the U.S. Previous studies indicated that this pathogen, although capable of causing disease in the absence of other pathogens in pathogenicity trials, is an opportunistic pathogen of oak trees, and acting in concert with other biotic and abiotic factors. Through the results of this study, we sought to find the geographic distribution of *Diplodia* spp. in the Mid-Atlantic region and observe relationships among isolates from Europe and the U.S. This survey showed that *Diplodia* spp. are more geographically widespread and associated with more oak hosts than previously reported. Although capable of causing disease, *Diplodia* spp. were not observed as the inciting agent of ongoing large-scale mortality events in the sampled areas, although their role in previous mortality events at these sites could not be assessed. This study revealed limited genetic diversity among *Dc* isolates from different geographic locations which may indicate *Dc* is a cosmopolitan pathogen, present from oak on both continents. However, secondary introductions are still possible and may reflect some of the fine structure observed in both the EF-1α tree and the 3-gene phylogeny. The occurrence of *Diplodia* outbreaks in geographically isolated regions, especially in more vulnerable coastal locations, may indicate that trees in these regions are more predisposed to secondary opportunistic pathogens. This study broadens the understanding of *Diplodia* distribution and its ecology, with evidence supporting *Dc* as a native pathogen to this region.

## Acknowledgments

Thanks to Isabel Munck, Jordi Luque, Shannon Lynch, and Srđan Aćimović for providing *Diplodia* isolates. Special thanks to Blake Ferreira for his field assistance. Funding for S.L.F. was provided by USDA Forest Service, Northeastern Area, State and Private Forestry (grant number 18-DG-11420004-121) and M.T.K. by funds from West Virginia Agricultural and Forestry Experiment Station, Morgantown, WV. Scientific article No. XXXX of the West Virginia Agriculture and Forestry Experiment Station, Morgantown, West Virginia, USA, 26506.

